# Model-based assessments of differential introgression and linked natural selection during divergence and speciation

**DOI:** 10.1101/786038

**Authors:** Arun Sethuraman, Vitor Sousa, Jody Hey

## Abstract

Demographic changes such as fluctuating population size and differential introgression can mask the effects of natural selection, and affect rates of genome evolution, local adaptation, reproductive isolation, and eventual speciation. Besides identifying differentially introgressing genes (and genomic regions) that are “labeled” to be retroactively causal to adaptive evolution and speciation, there is significant impetus to understand, and perhaps estimate the underlying demography that affects current genomic diversity. Using model-based likelihood methods to directly estimate, and decouple the effects of differential intogression and demography across genomic loci offers an ideal solution to detect differential introgression, and population demography and build hypotheses around its underlying evolutionary processes. We describe a computationally efficient parallelized implementation of mixture-model based isolation with migration (IM) analyses to assign loci to classes based on their shared coalescent histories (population sizes, or migration rates). We apply this method to several genomic data sets (great apes - chimpanzees and bonobos, Anopheles mosquitoes, threespine sticklebacks, Mullerian mimics of Heliconius butterflies, mice, European rabbits, and fruitflies), that have been previously characterized (perhaps erroneously) using genome-wide scans of differentiation. We show that we cannot reject a model of differential introgression, or linked selection across a majority of species analyzed, with two species showing the combined effects of differential introgression and linked natural selection across multiple, non-independent genomic loci.

---

In recent years, a number of studies have reported findings of the hidden hand of natural selection in contributing to species formation, as revealed by signals differential introgression in which some parts of genomes are exchanged between diverging populations while others are not. (e.g. Duranton et al. [8], Feulner et al. [10], Geraldes et al. [13], Kronforst et al. [16], Martin et al. [18], Ravinet et al. [24], Roux et al. [25], Tavares et al. [33], Turner and Hahn [34], Turner et al. [35]). Although the evidence is indirect, the argument is straightforward; for if it is true that only some portions of the genome are exchanged between diverging populations, then it follows that some selective force is preventing exchange at complementary portions of the genome (see Ravinet et al. [24], Roux et al. [25], Sousa and Hey [28]). A process of speciation that includes natural selection against introgression, is one in which selection at the genic level is acting indirectly as a diversifying force at the species level, and it is a far more complex process than one of simple allopatric divergence. Consequently the many reports of differential introgression during divergence have been used to support the case that our overall understanding of the speciation process is changing in important ways Abbott et al. [1], Feder et al. [9], Seehausen et al. [26], Sousa and Hey [28].

A potential difficulty with interpretations of differential introgression lies in the fact that such conclusions are themselves indirect, as they are usually based upon static patterns of variation observed within and between populations. A typical approach is to perform a genome scan, measuring divergence in windows along the genome, and identifying regions of markedly low or high divergence Cruickshank and Hahn [7], Payseur and Rieseberg [22]. The difficulty in this context is that divergence, by any measure, can be affected by other evolutionary factors besides gene flow. In particular, as some have pointed out Cruickshank and Hahn [7], Noor and Bennett [21], variation in divergence can arise even in the absence of gene exchange as a byproduct of natural selection within populations. For example, genetic divergence between populations is frequently measured using Wright’s *F*_*ST*_ Wright [40, 41], which is the proportion of variation that lies between populations. However, as a proportion, *F*_*ST*_ can be low either because the ‘between’ portion of the variation is low or because the ‘within’ portion is high. Loci that experience forces that lessen variation within populations, but do not otherwise affect divergence, will exhibit a high *F*_*ST*_, as will loci that have typical polymorphism within populations, but also have an elevated level of divergence. A simpler measure of divergence is the average number of base pair differences between sequences from different populations (frequently referred to as *d*_*XY*_ Nei and Miller [19]). Although not a proportion, this measure includes both variation within and between populations and can be elevated or reduced because of changes in either.

Our approach is to adapt a demographic model-based framework to allows for variation in the demographic parameters across the genome, in a way that it can account for the effects of natural selection on a subset of loci in the analysis Barton and Bengtsson [2], Charlesworth [5], Galtier et al. [11], Petry [23], Storz et al. [30]. We use an isolation-with-migration model (Fig. 1) which includes parameters for genetic drift (i.e. effective population size, gene flow, and time of population separation Latter [17], Nielsen and Wakeley [20], Takahata [32], Wakeley and Hey [38]). We extend the method of Sousa et al. [29] that allows for multiple sets of demographic parameters in a mixture model in which loci are grouped on the basis of their inferred demographic parameters, to deal with genome-wide data and perform likelihood ratio tests to infer if variation in introgression is required to explain the data. To improve computational efficiency, we developed a method that distributes the proposed Metropolis Coupled Markov Chain Monte Carlo (MC3) method across processors, providing a nearly linear speed-up in computational times, thus making it possible to perform inference using larger data sets (of the order of hundreds of loci) than previously feasible. This is implemented in the program `MigSelect`. Analyses of genomic data from seven paradigmatic species pairs across a variety of taxa under this framework reveals evidence of differential introgression and linked selection effects across several species, including new discoveries of differential introgression between species of chimpanzees (*Pan troglodytes verus* and *Pan troglodytes troglodytes*).

**Figure 1:**
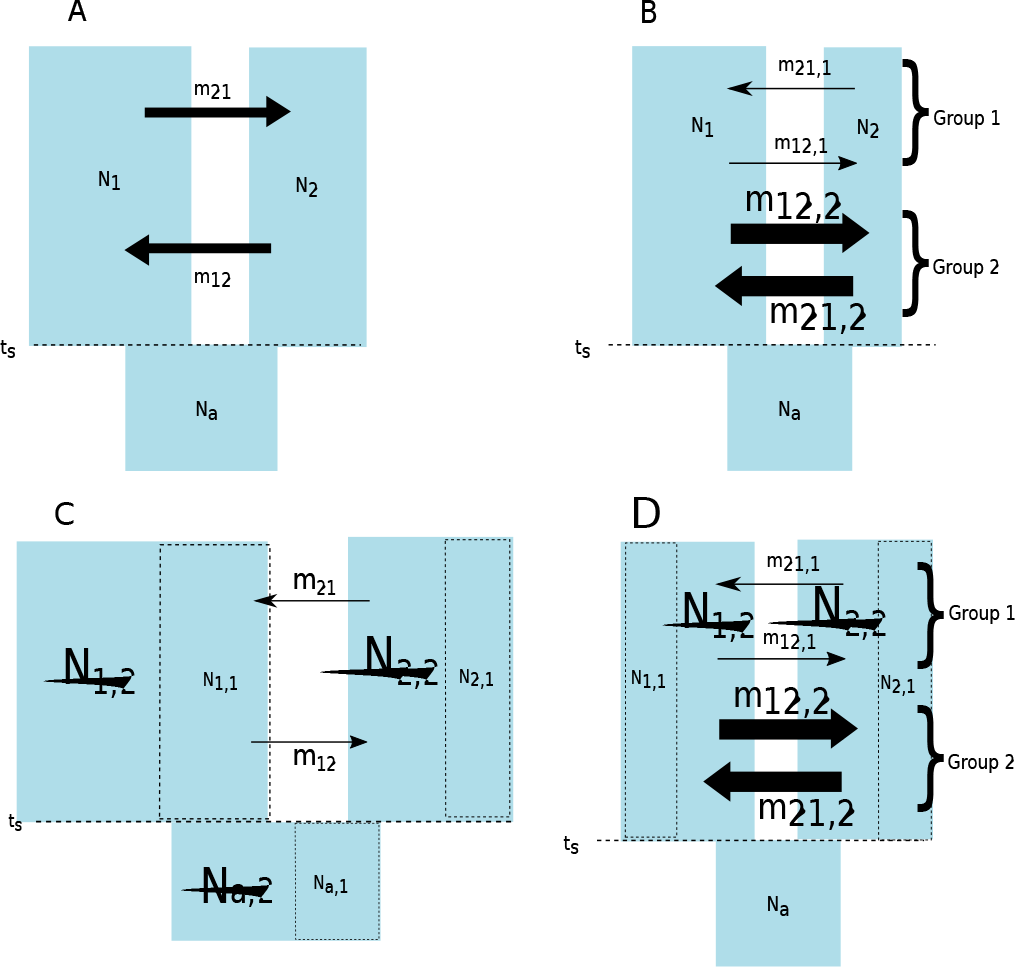
(A) Depiction of a typical Isolation with Migration (IM) model of Nielsen and Wakeley [20], which describes an ancestral population of size *N*_*a*_ splitting into two populations of sizes *N*_1_ and *N*_2_ at time *t*_*s*_. The two populations subsequently continue to exchange genes at rates *m*_12_ and *m*_21_ from population 1 into 2 and vice versa respectively. (B) This model describes a scenario where there are two groups of loci, 1 and 2, with different migration rate distributions (*m*_12,1_,*m*_21,1_,*m*_12,2_,*m*_21,2_). Note that this model only contains one group of effective population sizes, *N*_1_,*N*_2_, and *N*_*a*_. (C) This depicts a scenario where there are two groups loci with different effective population size distributions, 1 and 2 (*N*_1,1_,*N*_2,1_,*N*_*a*,1_ and *N*_1,2_,*N*_2,2_,*N*_*a*,2_). This model also only parametrizes one group of migration rates, *m*_12_ and *m*_21_. A full model which nests all models (A,B, and C) would in turn have two groups of loci for both effective population sizes, and migration rates, and is shown in panel D

## Results

### Inference Framework

Following the framework developed by Sousa et al. [29], we propose a series of three tests that together can decouple the effects of differential introgression and linked selection across loci. For a given data set the process begins by sampling from an MCMC simulation under a full IM model in which there are two sets of population size parameters *and* two sets of migration parameters (see Fig. 1). Then the sampled genealogies are used to construct a function that is an estimate of the joint posterior density for all of these parameters Hey and Nielsen [15]. Following Sousa et al. [29] we obtained the joint parameter estimate by optimizing this function while effectively integrating over both assignment vectors (i.e. for population size parameter sets, and for migration rate parameter sets). Furthermore, under a uniform prior on demographic parameters, this joint posterior density is proportional to the likelihood, and the estimated surface can be used for log likelihood ratio (LLR) tests of nested models. Specifically, we develop three LLR tests - (1) Test of Differential Introgression, wherein there are two sets of loci with estimably different migration rates (2) Test of Linked Selection, wherein there are two sets of loci with different effective population sizes, and (3) Test of Independence of Loci under differential introgression versus linked selection effects.

### Choice of Species Pairs

We analyzed genomic data from a variety of taxa (see Table 1), exhibiting several interesting natural history characteristics, including naturally non-hybridizing species (e.g. *P. paniscus verus* and *Pan troglodytes troglodytes*), species that freely hybridize in the wild (eg. *Anopheles gambiae* - *A. coluzzii*, formerly M subtype, and *A. gambiae sensu stricto*, formerly S subtype), and species that exhibit some levels of noted hybrid incompatibility (eg. *Mus musculus* - hybrid males showing lower fertility, or inviability). Species analyzed also form complexes under a variety of ecological speciation mechanisms - allopatry (eg. *P.p.verus*,*P.t.troglodytes*, and parapatry (e.g. *O.c.cuniculus*,*O.c.algirus*, *Gasterosteus* river versus lake forms). Ten replicate runs of `IMa2p` (Sethuraman and Hey [27]) and `MigSelect` were performed, discarding 100, 000 MCMC iterations as burn-in, and sampling 10, 000 ge-nealogies each. The 100, 000 genealogies sampled thus across replicate runs were then pooled and used in computing the marginal posterior density distributions of population mutation rates, migration rates, and divergence times, and in Likelihood Ratio Tests (LRT’s) of linked selection, differential introgression, and a test of independence of loci in nested models.

**Table 1:** Species pairs analyzed using `MigSelect`. Where genomic data was available, 200 random intergenic, unlinked, non-recombinant loci were sampled based on binned 10 kbps *F*_*ST*_ windows across the genome. Also shown are the ranges of pairwise *F*_*ST*_ values, computed across windows using the estimator of Weir and Cockerham (1984). Non-genomic data-sets were curated from GenBank, or Dryad, aligned, pruned, phased, and used in IM analyses.

| Species | Pop 1 | Pop 2 | Loci | Citation | $F_{st}^{Low}$ | $F_{st}^{High}$ |
| --- | --- | --- | --- | --- | --- | --- |
| *Oryctolagus cuniculus* | *cuniculus* | *algirus* | 44 | Sousa et al. 2013 | -0.04 | 0.92 |
| *Anopheles coluzzii* | *coluzzii* | *gambiae* | 36 | Turner and Hahn 2007, Turner et al. 2005 | 0.02 | 0.92 |
| *Gasterosteus aculeatus* - Germany | Lake | Riverine | 200 | Feulner et al. 2015 | -0.08 | 0.73 |
| *Heliconius melpomene* | *aglaope* | *amaryllis* | 200 | Supple et al. 2013 | -0.12 | 0.95 |
| *Mus musculus* | *musculus* | *domesticus* | 14 | Harr 2006 | 0.18 | 0.95 |
| *Pan* | *paniscus* | *trogodytes*<br>*trogodytes* | 200 | Prado-Martinez et al. 2013 | -0.09 | 0.46 |
| *Pan* | *trogodytes*<br>*trogodytes* | *trogodytes*<br>*verus* | 200 | Prado-Martinez et al. 2013 | -0.09 | 0.82 |
| *Drosophila pseudoobscura* | *pseudoobscura* | *persimilis* | 200 | Machado, Kingan and Garigan (unpublished) | -0.04 | 0.85 |

### Tests of Significant Migration

Tests of significant migration under two species IM models using `IMa2p` could not reject the null hypothesis of no significant migration in six out of the eight species pairs analyzed (Table 2). Out of these, sampled loci across *Heliconius* butterflies, *Oryctolagus* rabbits, and *Anopheles* mosquitoes showed signatures of significant bidirectional migration across all loci, whereas the great ape species pairs (*P.paniscus* vs *P.t.troglodytes*, and *P.t.troglodytes* vs *P.t.verus*), and *D. pseudoobscura* vs *D. persimilis* estimate significant unidirectional gene flow. Interestingly, despite previous reports of significant migration between natural populations (see Feulner et al. [10], Harr [14]), our `IMa2p` analyses do not estimate any significant migration between these species pairs.

**Table 2:** LLR Tests of significant migration between sampled populations using `IMa2p`, under the *H*_0_ that there is no significant migration between sampled populations, and *H*_*A*_ that there is significant migration (adjudged at a P-value of 0.05). Significant migrations are shown in red.

| Species | $m_{0 \rightarrow 1}$(LLR) | $m_{1 \rightarrow 0}$(LLR) | Decision |
| --- | --- | --- | --- |
| *Heliconius melpomene* vs *aglaope amaryllis* | 0.29(p<0.001) | 0.48(p <0.001) | Reject $H_0$ |
| *Oryctolagus cuniculus cuniculus* vs *algius* | 0.37(p < 0.001) | 0.36(p < 0.001) | Reject $H_0$ |
| *Anopheles coluzzii* vs *gambiae* | 9.944(p<0.001) | 5.419(p<0.001) | Reject $H_0$ |
| *Pan paniscus* vs *trogodytes troglodytes* | 0.129(ns) | 0.178(p < 0.001) | Reject $H_0$ for $m_{1 \rightarrow 0}$ (Bonobo→Central) |
| *Pan troglodytes troglodytes* vs *trogodytes verus* | 0.000(ns) | 2.05 (p < 0.001) | Reject $H_0$ for $m_{1 \rightarrow 0}$ (Central→Western) |
| *Drosophila pseudoobscura* vs *persimilis* | 641.585(p<0.001) | 0.000(ns) | Reject $H_0$ for $m_{0 \rightarrow 1}$ (pers.→pseudo.) |
| *Gasterosteus aculeatus* Germany Lake vs Riverine | 0.06(ns) | 0.005(ns) | Cannot reject $H_0$ |
| *Mus musculus musculus* vs *domesticus* | 0.000(ns) | 0.17(ns) | Cannot reject $H_0$ |

### Tests of Differential Introgression

Five out of eight species pairs analyzed using `MigSelect` failed to reject the null hypothesis of no differential introgression effects across their genomes at a significance level of 0.01 (Table 3).

**Table 3:**
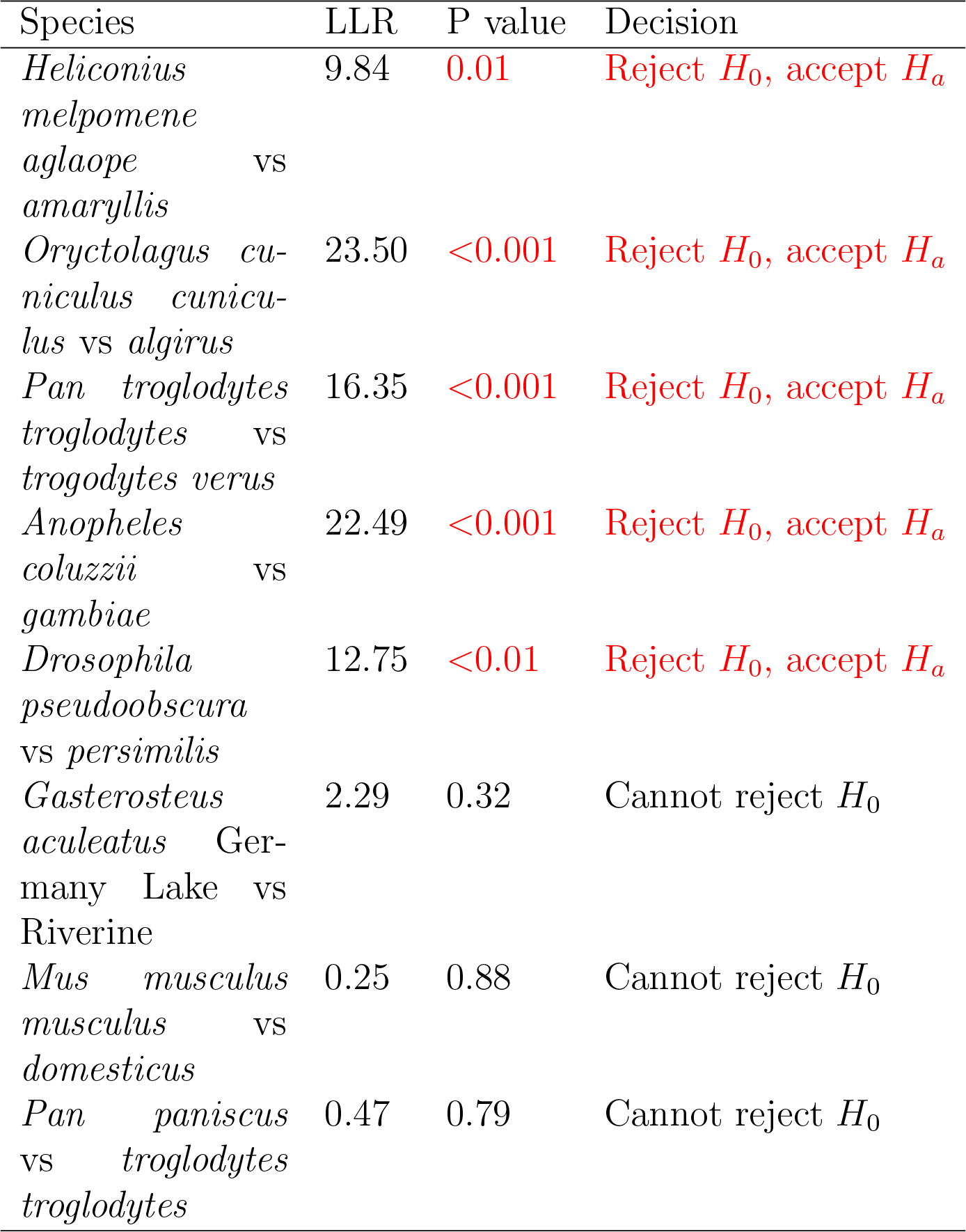
Tests of differential introgression using two groups of loci for migration rates, using the *H*_0_ that there is only one group of loci for migration rates versus the *H*_*A*_ that there are two groups of loci. Significance was adjudged at a P-value of 0.05, and significant tests are indicated in red.

Analyses of the functions of loci enriched for ‘low’ and ‘high’ migration rates in each species pair discovered genes that encoded for LIM binding proteins, sodium ion channel proteins, 3L meiotic protein, and AGAP010317, AGAP010315, which have all been reported to be present on predicted divergence islands (White et al. [39]) in *Anopheles*.

Similar analyses of functional genes in sticklebacks showed that loci classified as having ‘low’ migration rates include large regions between 193kb - 216kb, 281kb - 302kb (including the *slu7*,*kinesin*, *gpcr* genes), 313kb - 379kb (including the highest divergence interval, as identified by Supple et al. [31]), 405kb - 455kb (including the *optix* genic region, which was also previously identified as containing high divergence in the Peruvian Hybrid Zone of *H. melpomene*), and a small 1kb region between 553kb - 554kb of the B/D interval around the *optix* gene (see Fig. 4 of Supple et al. [31]). Our analyses also classified a few loci as displaying ‘high’ migration rates. These include a large region between 16kb - 27kb, a 1kb region between 92kb - 93kb, a large region between 454kb - 532kb, and 555kb - 560kb. The latter two regions are immediately downstream of regions that contain loci that are enriched for “low” migration around the *optix* gene - potentially containing regulatory elements, congruent with the hypothesis of natural selection on color pattern variation in these hybrid zones.

Functional annotation enrichment analyses of loci that are putatively under differential introgression effects in great ape species pairs (79 out of 200 loci in *P. paniscus* vs *P. troglodytes*) discovered several ‘hits’ to GO terms mapped to the *hg38* human genome annotation (see Supplemental Table:…). In *D. pseudoobscura* vs *D. persimilis*, most loci classified as having ‘high’ migration rates occurred around a chromosome 2 inversion, suggesting support for gene flow for loci within this inversion, as also reported by Machado et al. 2007.

### Tests of Linked Selection

Tests of linked natural selection also failed to reject the null hypothesis of no linked selection effects (i.e. two different groups of loci with different effective population sizes) in six of eight species pairs analyzed (Table 4) at a significance level of 0.01.

**Table 4:** Tests of linked selection effects using two groups of loci for effective population sizes using the *H*_0_ that there is only one group of loci, versus *H*_*A*_ that there are two groups of loci for effective population sizes. Significance was adjudged at a P-value of 0.05. Species pairs that are significant for tests of linked selection effects are shown in color (red - indicates species that were not significant for tests of differential introgression, blue - species that were significant for tests of differential introgression.

| Species | LLR | P value | Decision |
| --- | --- | --- | --- |
| *Gasterosteus aculeatus* Germany Lake vs Riverine | 15.92 | <0.01 | Reject $H_0$, accept $H_a$ |
| *Mus musculus musculus* vs *domesticus* | 13.64 | <0.01 | Reject $H_0$, accept $H_a$ |
| *Pan paniscus* vs *troglodytes troglodytes* | 241.50 | <0.001 | Reject $H_0$, accept $H_a$ |
| *Oryctolagus cuniculus cuniculus* vs *algius* | 128.50 | <0.001 | Reject $H_0$, accept $H_a$ |
| *Anopheles coluzzii* vs *gambiae* | 14.25 | <0.01 | Reject $H_0$, accept $H_a$ |
| *Drosophila pseudoobscura* vs *persimilis* | 1099.00 | <0.001 | Reject $H_0$, accept $H_a$ |
| *Heliconius melpomene aglaope* vs *amaryllis* | 7.06 | 0.07 | Cannot reject $H_0$ |
| *Pan troglodytes troglodytes* vs *trogodytes verus* | 5.78 | 0.12 | Cannot reject $H_0$ |

Across *Oryctolagus* species pair loci, `MigSelect` analyses show a clear differentiation of loci into three classes of loci - those with high population size, and high migration rate (A31-GBE,A37-EXT1,A38-LUM), loci with high migration rate, and low population sizes (X3-MAOA,X16-DIAPH2), and with low migration rates, and low population sizes (X23-SHOX,X19-G6PD,X18-FMR1,X15-AMOT,X17-F9,A43-STAG1), consistent with the observations of Carneiro et al. [4] and Sousa et al. [29].

Analyses of linked selection effects in *Gasterosteus* species pair loci discovered 20 out of the 200 loci to be classified as having ‘higher’ effective population size than others (see Supplemental Table …). Among these, was Locus2 (GroupI:2150001-2160000), which is about 100kb upstream of the *ATP1a1* gene, which is known to be gene involved in osmotic regulation (and previously identified as an *F*_*ST*_ outlier by Feulner et al. [10] across all populations analyzed. As reported by Feulner et al. [10], the *Eda* gene or its immediate neighboring regions were not enhanced for linked selection effects. Also of note are two nearly consecutive regions on linkage groupXX(17730001-17740000,17760001-17770000) which are also classified as being under the effects of linked natural selection effects. These regions contain exons belonging to the *Lin28* and *MTF1* genes, which are implicated in *IGF2* translational regulation, and in metal homeostasis respectively.

In the *Mus musculus* species pair, seven out of the fourteen loci analyzed (*Ppp1r3b*, *Sec63*, *Nr1d2*, *Wdt2*, *Bnc1*, *XP 620246*, *Ggh*, *Melk*) were classified as loci with higher effective population sizes, than the others. These loci had a wide range of variability in estimated *F*_*st*_ (0.13 (*Ggh*)−0.72(*Sec63*) (see Supplementary Table 1 of Cruickshank and Hahn [7]), further exhibiting the ‘confounding’ effects of demography on estimates of differentiation or divergence.

Analyses of *P.t.verus* versus *P.t.troglodytes*, which estimated signatures of linked natural selection effects identified 14 loci (out of 200) that were classified to have greater effective sizes than others, and thus potentially under the effects of linked natural selection. Identifiable human orthologs to these ‘outlier’ loci include those involved in G-protein receptor functions, fukutin, and membrane protein synthesis.

In the *Drosophila* species pair, loci classified as being enriched for ‘low’ effective population size occurring around a chromosome 2 inversion (9Mb-17Mb) as reported by Machado et al. 2007 and McGaugh and Noor 2012. Additional loci with ‘low’ effective sizes were also identified on chromosomes 3 and 4.

### Tests of Independence of Genomic Loci

A Fisher’s exact test of independence between loci that were classified as under differential introgression effects versus those under linked selection effects failed to reject the null hypothesis of independence in two species pairs - *Anopheles* and *Drosophila* (Table 5). In the *Anopheles* complex, one locus (cytochrome P450 gene) on the X chromosome was determined to enriched for low population size, and high migration rate, indicative of possible adaptive introgression on the X chromosome (also shown to have low differentiation - *π* < 0.001 by Turner et al. [35]).

**Table 5:** Fisher’s exact tests of independence of loci that are affected by both differential introgression and linked selection effects. Results are thus only shown for species pairs that were significant for two groups of analyzed loci for migration, and effective population sizes. Significance was adjudged at a P-value of 0.05, and indicated in red.

| Species | P value | Decision |
| --- | --- | --- |
| *Anopheles coluzzii* vs *gambiae* | <0.001 | Reject $H_0$ |
| *Drosophila pseudoobscura* vs *persimilis* | 0.04 | Reject $H_0$ |
| *Oryctolagus cuniculus* vs *algericus* | 0.06 | Cannot reject $H_0$ |

In the *Drosophila* species complex, we see a consistent overlap between loci that are enriched for ‘high’ migration rates and those with ‘low’ effective population size around a chromosome 2 inversion, suggesting support for gene flow for loci within this inversion, as also reported by Machado et al. 2007. Loci classified as having ‘low’ effective sizes on chromosomes 3 and 4 were also enhanced for ‘high’ migration rates. Several loci on chromosome 4 (group 3) were enhanced for ‘high’ effective sizes, which contained loci overlapping with having been enhanced for ‘low’ migration rates. We hypothesize that these loci are potentially under the effects of balancing selection in each species, and could potentially represent speciation islands.

## Discussion

Adaptive evolutionary history post-isolation is often complicated by the effects of differential migration and linked natural selection between diverged populations of a species. Traditional methods of quantifying diversity and differentiation across genomic regions, including genome-wide scans of *F*_*ST*_, *d*_*XY*_, and similar statistics fail to capture these differences. As pointed out by Cruickshank and Hahn [7], all measures of divergence and differentiation can be affected by other evolutionary factors besides gene flow. For example, background selection, or negative selection on deleterious alleles (perhaps from the migrant population) at genomic loci can result in reduced absolute or relative diversity within populations, and thus increased divergence or differentiation between populations (Charlesworth et al. [6],Noor and Bennett [21],Cruickshank and Hahn [7]). Similarly, positive selection and selective sweeps can cause reduction in absolute or relative genomic diversity at linked neutral sites within populations, and increased relative divergence between populations (Via [36], Via and West [37]). There is as yet no consensus on which measure (relative or absolute) is more powerful in detecting loci experiencing either differential migration or linked selection effects (Cruick-shank and Hahn [7],Geneva et al. [12]). Numerous lines of evidence also point to the role of recombination in maintenance of genomic diversity in the face of gene flow post-divergence (Noor and Bennett [21], Burri et al. [3]). Regions of low recombination (eg. pericentromeric, or telomeric regions) can harbor low genomic diversity due to hitchhiking, or background selection, thus portending greater detected differentiation and divergence between populations.Similarly, regions of chromosomal rearrangements exhibit lower diversity owing to incomplete lineage sorting of rearranged sections (Noor and Bennett [21]). It is thus difficult to identify the cause of variation in genomic diversity or divergence using a genome scan of summary statistics.

We present a model-based approach and a series of tests of linked selection, differential introgression, and a test of independence between the two effects to help tease these effects apart. We demonstrate its utility by analyzing simulated and empirical genomic data sets of varying sizes and natural histories. Implemented using a parallelized framework (using `OpenMPI`), `MigSelect` is fast (data not shown, but see Sethuraman and Hey [27], and scales easily to hundreds of genomic loci with computing power. The software is now freely available for academic use at www.github.com/arunsethuraman/MigSelect/.

## Methods

### Test of Linked Selection Effects

If directional selection (positive or negative) is systematically shaping polymorphism and divergence patterns at some loci, then these loci will appear to have altered effective population sizes, relative to loci that are not similarly affected, and sampled assignment vectors for population size parameters will tend to show a grouping of these loci. Finally a comparison of log likelihoods, for the full model (that includes two sets of population size parameters) against the nested model in which all loci share the same population size parameters, is expected to reveal a large difference.

We denote the full, or alternate model, *H*_*a*_, as the full set of parameters: 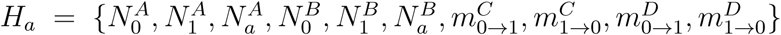, where *A* and *B* are the two sets of population size parameters, and *C* and *D* are the two sets of migration rate parameters. And we denote the null model *H*_0_ as having just a single set of population size parameters: 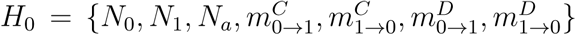. Then from a single set of genealogies sampled under the full model, we calculate the LLR statistic, *L*(*X*), as twice the difference between maximized log values from two joint density functions.

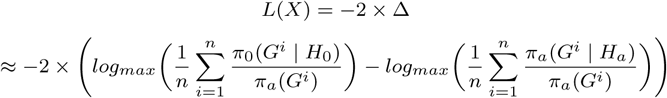

Here *G*^*i*^ is the *i*^*th*^ genealogy sampled from the posterior, *G*^*i*^ ∼ *P* (*X*|*H*_*a*_), and *π*_*a*_ denotes a prior probability calculated by analytic integration over the demographic parameters in *H*_*a*_ (Hey and Nielsen [15]).

### Test of Differential Introgression

If the fate of genes that introgress from one population to another, varies systematically, such that some portions of the genome experience different effective rates of gene flow than others, then we expect to observe a large difference in log likelihoods for models that do, and do not, include multiple sets of migration parameters. The test of differential introgression is very similar to that for linked selection effects, with the full model, *H*_*a*_, being the same. However in the case of differential introgression the null model includes only one set of migration parameters: 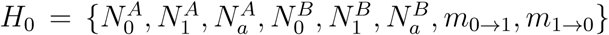, where *A* and *B* are the two sets of population size parameters. The LLR statistic *L*(*x*) is then similarly computed using Eqn..

### Test of independence of loci affected by differential introgression and linked selection effects

We consider a third test, which determines if there are signatures of evolutionary processes that simultaneously affect population sizes, and migration rates (and hence linked selection, and differential introgression effects) between sampled populations. Consider the null hypothesis (*H*_0_) that the groups of loci that are differentially introgressing are independent of loci that exhibit linked selection effects. Alternately, consider the *H*_*a*_ that loci that are differentially introgressing and the loci that have linked selection effects are non-independent. We use a Fisher’s exact test of independence with contingency tables of assignments of loci, computed across all sampled assignment vectors in the method of Sousa et al. [29].

### Empirical Datasets

Data from all previous studies that utilized targeted sequencing (< 100 loci) were aligned, phased, pruned to exclude recombinant, and invariant segments, prior to demographic analyses. In data sets where whole-genomic data was available, variant calls were extracted, genome-wide estimates of relative differentiation (*F*_*ST*_) were computed, filtered by genotyping/sequencing quality, and loci sampled uniformly from across this distribution by ranking *F*_*ST*_ values. Sampled loci were then subject to the same IM model assumption checks as before, and for most pairs of species, 200 representative loci were used in tests of linked selection and differential introgression.

All data sets, and scripts to produce them have been deposited with Dryad (DOI:…).

## Acknowledgments

This work was supported by NSF Grant:1564659 to AS and JH, NIH Grant:R01GM078204 to JH.

